# ArrowSAM: In-Memory Genomics Data Processing Using Apache Arrow

**DOI:** 10.1101/741843

**Authors:** Tanveer Ahmad, Nauman Ahmed, Johan Peltenburg, Zaid Al-Ars

**Author notes:** {, }.

## Abstract

The rapidly growing size of genomics data bases, driven by advances in sequencing technologies, demands fast and cost-effective processing. However, processing this data creates many challenges, particularly in selecting appropriate algorithms and computing platforms. Computing systems need data closer to the processor for fast processing. Traditionally, due to cost, volatility and other physical constraints of DRAM, it was not feasible to place large amounts of working data sets in memory. However, new emerging storage class memories allow storing and processing big data closer to the processor. In this work, we show how the commonly used genomics data format, Sequence Alignment/Map (SAM), can be presented in the Apache Arrow in-memory data representation to benefit of in-memory processing and to ensure better scalability through shared memory objects, by avoiding large (de)-serialization overheads in cross-language interoperability. To demonstrate the benefits of such a system, we propose ArrowSAM, an in-memory SAM format that uses the Apache Arrow framework, and integrate it into genome pre-processing pipelines including BWA-MEM, Picard and Sambamba. Results show 15x and 2.4x speedups as compared to Picard and Sambamba, respectively. The code and scripts for running all workflows are freely available at https://github.com/abs-tudelft/ArrowSAM.

## I. Introduction

Genomics is projected to generate the largest big data sets globally, which requires modifying existing tools to take advantage of new developments in big data analytics and new memory technologies to ensure better performance and high throughput. In addition, new applications using DNA data are becoming ever more complex, such as the study of large sets of complex genomics events like gene isoform reconstruction and sequencing large numbers of individuals with the aim of fully characterizing genomes at high resolution [1]. This underscores the need for efficient and cost effective DNA analysis infrastructures.

At the same time, genomics is a young field. To process and analyze genomics data, the research community is actively working to develop new, efficient and optimized algorithms, techniques and tools, usually programmed in a variety of languages, such as C, Java or Python. These tools share common characteristics that impose limitations on the performance achievable by the genomics pipelines.

- These tools are developed to use traditional I/O file systems which incur a huge I/O bottleneck in computation due to disk bandwidth [2]. Each tool reads from the I/O disks, computes and writes back to disk.
- Due to the virtualized nature of some popular languages used to develop genomics tools (such as Java), these tools are not well suited to exploit modern hardware features like Single-instruction multiple-data (SIMD) vectorization and accelerators (GPU or FPGAs).

This paper proposes a new approach for representing genomics data sets, based on recent developments in big data analytics to improve the performance and efficiency of genomics pipelines. Our approach consists of the following main contributions:

- We propose an in-memory SAM data representation, called ArrowSAM, created in Apache Arrow to place genome data in RecordBatches of immutable shared memory objects for inter-process communication. We use DRAM for ArrowSAM placement and inter-process access.
- We adapt existing widely-used genomics data preprocessing applications (for alignment, sorting and duplicates removal) to use the Apache Arrow framework and to benefit from immutable shared memory plasma objects in inter process communication.
- We compare various workflows for genome preprocessing, using different techniques for in-memory data communication and placement (for intermediate applications), and show that ArrowSAM in-memory columnar data representation outperforms.

The rest of this paper is organized as follows. Section II discusses background information on genomics tools and Apache Arrow big data format. Section III presents the new ArrowSAM genomics data format. Section IV shows how to integrate Apache Arrow into existing genomics tools, while Section V discusses the measurement results of these new tools. Section VI presents related work in the field. Finally, Section VII ends with the conclusions.

## II. Background

This section provides a short description of DNA sequence data pre-processing tools, followed by a brief introduction to the Apache Arrow framework.

### A. DNA pre-processing

After DNA data is read by sequencing machines, alignment tools align reads to the different chromosomes of a reference genome and generate an output file in the SAM format. BWA-MEM [3] is a widely used tools for this purpose. The generated SAM file describes various aspects of the alignment result, such as map position and map quality. SAM is the most commonly used alignment/mapping format. To eliminate some systematic errors in the reads, some additional data preprocessing and cleaning steps are subsequently performed, like sorting the reads according to their chromosome name and position. Picard [4] and Sambamba [5] are some tools commonly used for such operations. This is followed by the mark duplicates step, where duplicate reads are removed by comparing the reads having the same map positions and orientation and selecting the read with the highest quality score. Duplicate reads are generated due to the wetlab procedure of creating multiple copies of DNA molecules to make sure there are enough samples of each molecule to facilitate the sequencing process. Again, Picard and Sambamba are commonly used here.

### B. Apache Arrow

To manage and process large data sets, many different big data frameworks have been created. Some examples include Apache Hadoop, Spark, Flink, MapReduce and Dask. These frameworks provide highly scalable, portable and programmable environments to improve storage and scalability of big data analytics pipelines. They are generally built on top of high-level language frameworks such as Java and Python to ensure ease of programmability. However, such high-level languages induce large processing overheads, forcing programmers to resort to low-level languages such as C to process specific computationally intensive parts of big data applications. On the other hand, heterogeneous components like FPGAs and GPUs are being increasingly used in cluster and cloud computing environments to improve performance of big data processing. These components are, however, programmed using very close-to-metal languages like C/C++ or even hardware-description languages. The multitude of technologies used often results in a highly heterogeneous system stack that can be hard to optimize for performance. However, combining processes programmed in different languages induces large inter-process communication overheads (so called data (de)serialization) whenever the data is moved between such processes.

To mitigate this problem, the Apache Arrow [6] project was initiated to provide an open standardized format and interfaces for tabular data in-memory. Using language-specific libraries, multiple languages can share in-memory data without any copying or serialization. This is done using the plasma inter-process communication component of Arrow, that handles shared memory pools across different processes in a pipeline [7].

## III. Arrowsam data format

This paper proposes a new in-memory genomics SAM format based on Apache Arrow. Such a representation can benefit from two aspects to improve overall system throughout: one is related to the tabular nature of genomics data and the other related to cross-language interoperability. Using Arrow ensures efficient genomics data placement in memory to gain maximum throughput and parallel data access efficiency. The tabular genomics data format can benefit from the standardized, cross-languages in-memory data representation of Arrow, that provides insight into the organization of the data sets through its schema.

In order to enable genomics applications to use Apache Arrow, two different contributions are needed. First, we need to define an Arrow in-memory representation of the corresponding genomics SAM data format. Second, the applications and tools using the data need to be adapted to access the new format as shown in Figure 1(a). In the following, these two contributions are discussed.

**Fig. 1.**
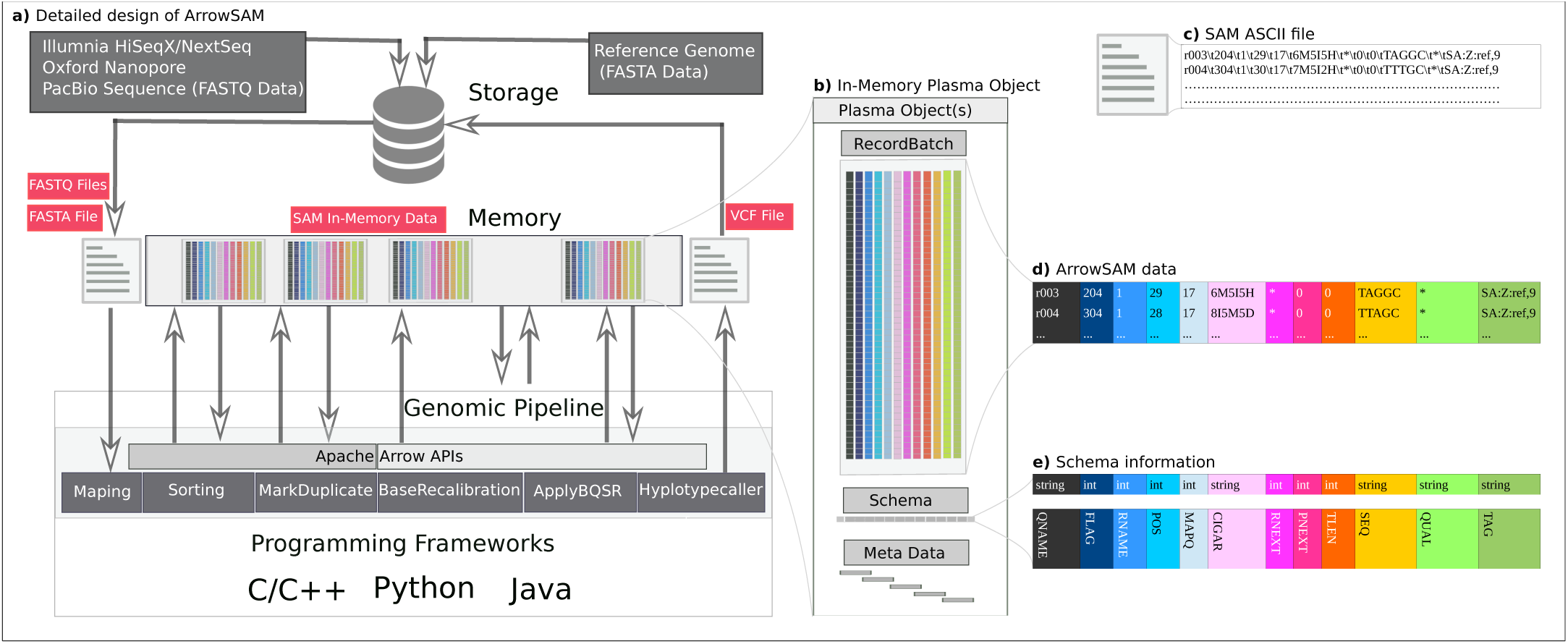
a) Genomics pipeline using ArrowSAM format for all intermediate steps to allow in-memory intermediate data storage, which means I/O disk access is only needed to load data into memory at the beginning and to write data back to disk at the end. b) Arrow RecordBatch enclosed in plasma object store. c) SAM file in ASCII text format. d) SAM file in RecordBatch format. e) Schema specifies the data types of ArrowSAM.

The SAM file format is an ASCII-based tab delimited text format to represent DNA sequence data as shown in Figure 1(c). Its in-memory SAM representation is a columnar format that consists of the same fields (columns) used in SAM to store the corresponding sequence mapping data as shown in Figure 1(d). The Arrow frameworks requires defining the data types for each field in a schema stored as part of the data object, as shown in Figure 1(e). The schema defines the ArrowSAM data format as listed more explicitly in Table I.

**TABLE I.** ArrowSAM Schema

| Index | Field | Datatype |
| --- | --- | --- |
| 0 | QNAME | String |
| 1 | FLAG | Int32 |
| 2 | RNAME | Int32 |
| 3 | POS | Int32 |
| 4 | MAPQ | Int32 |
| 5 | CIGAR | String |
| 6 | RNEXT | Int32 |
| 7 | PNEXT | Int32 |
| 8 | TLEN | Int32 |
| 9 | SEQ | String |
| 10 | QUAL | String |
| 11 | TAG | String |

Arrow stores the columnar data fields in contiguous memory chunks in so-called RecordBatches as shown in Figure 1(b). Each RecordBatch is a combination of a schema, which specifies the types of data fields of the ArrowSAM record, the data itself, in addition to some meta data.

## IV. Implementation

### A. BWA-MEM integration

`BWA-MEM` aligns the raw read sequences against a large reference genome such as that of a human. We used ArrowSAM to store the mapping data produced by `BWA-MEM` from query and reference genome files. We modified `BWA-MEM` to use Arrow libraries to write each chromosome (1-22, X, Y and M) sequence mapping data in a separate Arrow RecordBatch. At the end of the alignment process, all the RecordBatches are assigned to a shared memory pool of plasma objects. Each plasma object has its own identifications (objectID). Tools that need to use the data generated by `BWA-MEM` can access this data managed by plasma through zero-copy shared memory access [8]. Doing so enables other tools to access all shared RecordBatches in parallel.

### B. Sorting through pandas dataframes

Pandas is a powerful and easy to use Python library, which provides data structures, data cleaning and analysis tools. Dataframes is an in-memory data library that provides structures to store different types of data in tabular format to perform operations on the data in columns/rows. Any row in a dataframe can be accessed with its index, while a column can be accessed by its name. A column can also be a series in pandas. Using dataframes illustrates the powerful capabilities of in-memory data representation. First of all, dataframes is able to sort the chromosomes in parallel using pandas built-in sorting function with Python Arrow bindings (PyArrow) while accessing data residing in-memory, which takes place across two applications written in different languages (one in C and the other in Python). Secondly, tools like `Picard` and `Sambamba` are used to sort the SAM file according to the chromosome name and start positions of each chromosome. This type of sorting becomes computationally intensive when the whole SAM file needs to be parsed and sorted based on the values stored in only two fields of that file. This can be parallelized and made more efficient in our approach. Using pandas dataframes, sorting each individual chromosome is performed based on the start position of reads in that particular chromosome. All the RecordBatches are fed to pandas dataframes to sort all the chromosomes in parallel. After sorting, the sorted chromosomes RecordBatches are assigned to plasma shared memory again for subsequent applications to access.

### C. Picard MarkDuplicate integration

After sorting the chromosomes data by their coordinates, the duplicate reads with low quality should be removed. The `Picard MarkDuplicate` tool is considered as a benchmark for this task. This tool reads the SAM file two times, first when building the sorted read end lists and then removing marked duplicates from the file. To overcome this I/O overhead, we just read the data as ArrowSAM format in-memory once, accessing only five fields (`QNAME`, `FLAG`, `RNAME`, `POS`, `CIGAR` and `RNEXT`) needed to perform the MarkDuplicate operation. We modified `htsjdk` (a java API used in `Picard` and many other tools for managing I/O access of DNA sequencing data files) and `MarkDuplicate` to read data from all RecordBatches in parallel from plasma shared memory. Our implementation processes this data in parallel in `Picard` and writes back the updated `FLAG` field in ArrowSAM which sets duplicate bit. After finishing this process, the shared memory plasma objects are available for further variant calling processes for in-memory and parallel execution.

## V. Evaluation

This section evaluates the speedup and efficiency achieved using ArrowSAM for pre-processing of sequence data while mapping, sorting and marking duplicates against existing frameworks.

### A. Experimental setup

All the experiments and comparisons are performed on a dual socket Intel Xeon server with E5-2680 v4 CPU running at 2.4 GHz. A total of 192 GB of DDR4 DRAM with maximum of 76.8 GB/s bandwidth is available for whole system.

We use Illumina HiSeq generated NA12878 dataset of whole exome sequencing (WES) of human with 30x sequencing coverage with paired-end reads and a read length of 100 bps. Similarly for whole genome sequencing (WGS), we use Illumina HiSeq generated NA12878 dataset sample SRR622461 with sequencing coverage of 2x with paired-end reads and a read length of 100 bps. Human Genome Reference, Build 37 (GRCh37/hg19) is used as a reference genome for all workflows in our experiments for both WES and WGS.

The code and scripts for running all workflows is freely available at https://github.com/abs-tudelft/ArrowSAM. Tools and libraries and their version numbers used in our experiments are listed in Table II.

**TABLE II.** Tools and libraries used in the experimental setup

| Tools/APIs | Version |
| --- | --- |
| BWA-MEM [3] | 0.7.17 |
| Picard [4] | 2.18.14 |
| Sambamba [5] | v0.6.8 |
| elPrep [9] | v4.1.5 |
| Arrow C/C++/Java [6] | 0.11.0 |
| PyArrow [10] | 0.11.0 |
| Plasma object store [7] | 0.11.0 |

### B. Performance evaluation

In this section, we compare our approach with state-of-the-art tools and approaches used for pre-processing of genomics sequencing data. All speedups are compared for best performance scenarios.

#### 1) Picard

tools are considered as benchmarks in genome analysis pipelines, such as `Picard MarkDuplicate`. This tool was adapted to use our ArrowSAM in-memory format. The `MarkDuplicate` process is compute intensive but there is a significant amount of time (approximately 30%) spent in I/O operations. `Picard` uses `htsjdk` as a base library to read/write SAM files. We modified this tool from two perspectives:

- Instead of reading from and writing to files, it now reads/writes from in-memory RecordBatches, using only those fields/columns necessary for `MarkDuplicate` operations.
- `Picard` is single threaded. We changed it to be multi-threaded so that each thread can operate on a separate chromosome data set.

The first modification provides the benefit of using only the required fields to perform `MarkDuplicate` operations instead of parsing all the reads in a SAM files. As shown in Figure 2, our implementation gives 8x and 21x speedups on `Picard` sorting for genome and exome data sets, respectively. Similarly for `Picard MarkDuplicate`, we achieve 21x and 18x speed-ups on for genome and exome data sets, respectively.

**Fig. 2.**
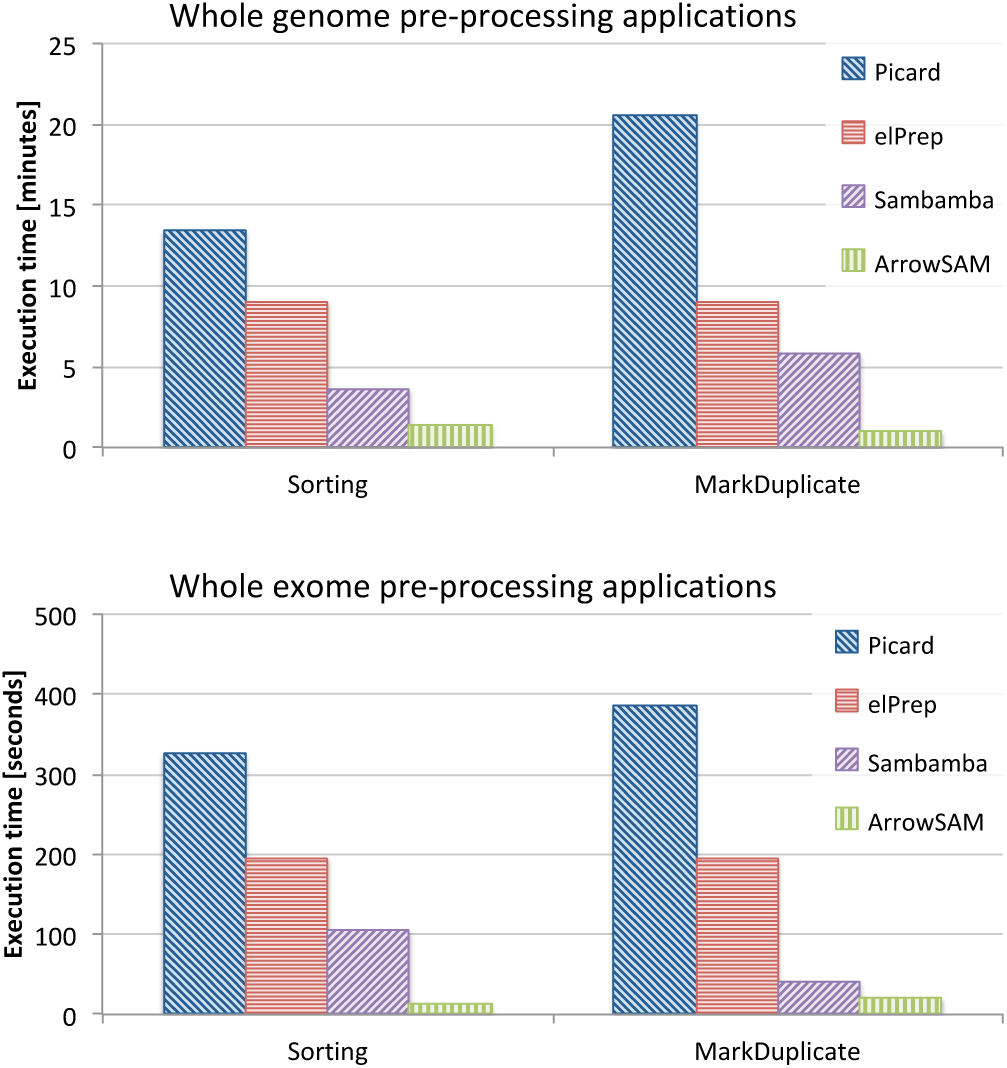
Execution time of Picard, Sambamba, elPrep and ArrowSAM based sorting and MarkDuplicate for whole genome (top) and whole exome (bottom) data sets.

#### 2) Sambamba

is a multi-threaded tool to manipulate SAM files for pre-processing steps in genomics pipelines. This tool gives close to linear speedup for up to 8 threads but adding more threads provides diminishing returns in performance on a multi-core system. The main reason behind this is the file system itself. The I/O communication gets saturated by initiating more threads and CPU performance also degrades because of cache contention [5]. As shown in Figure 2, our implementation gives 2x speedup on `Sambamba` sorting for both genome and exome data sets. Similarly for `Sambamba MarkDup`, we achieve 1.8x and 3x speedups for genome and exome data sets, respectively.

#### 3) elPrep

is the latest set of multi-threaded tools for pre-processing SAM files in-memory. We have also tested and compared these tools with our implementation for preprocessing applications and results show that our implementation gives more than 5x speedup over `elPrep`. `elPrep` performs sorting and mark duplicate in a single command, with a total run-time for both stages equally divided in run-time graphs Figure 2. Samblaster is yet another tool used for pre-processing SAM files, which is faster than `Sambamba`, but has not been considered for performance comparison here because it produces a different output for the `MarkDuplicate` stage than `Picard`.

### C. Discussion

#### 1) CPU usage

Figure 3 shows CPU utilization for standard `Picard` (left) as well as ArrowSAM-based (bottom) sorting and `MarkDuplicate` for whole exome data. In both sorting and duplicates removal stages, the parallelization offered by shared memory plasma objects results in a large speedup. All 25 chromosomes are sorted and duplicates are removed in parallel. In `Picard` sorting the CPU utilization is poor and the tool is mostly waiting for I/O. The average CPU utilization is only 5%. The `Picard MarkDuplicate` also has very low CPU utilization, although better than sorting. On the other hand, the CPU utilization of our implementation, which uses pandas dataframes is much better than `Picard` sorting. The CPU utilization of `MarkDuplicate` in our implementation remains close to 95% during the whole execution stage. The improved CPU utilization is due to Arrow in-memory storage and parallel execution of processes, each working on a different chromosome.

**Fig. 3.**
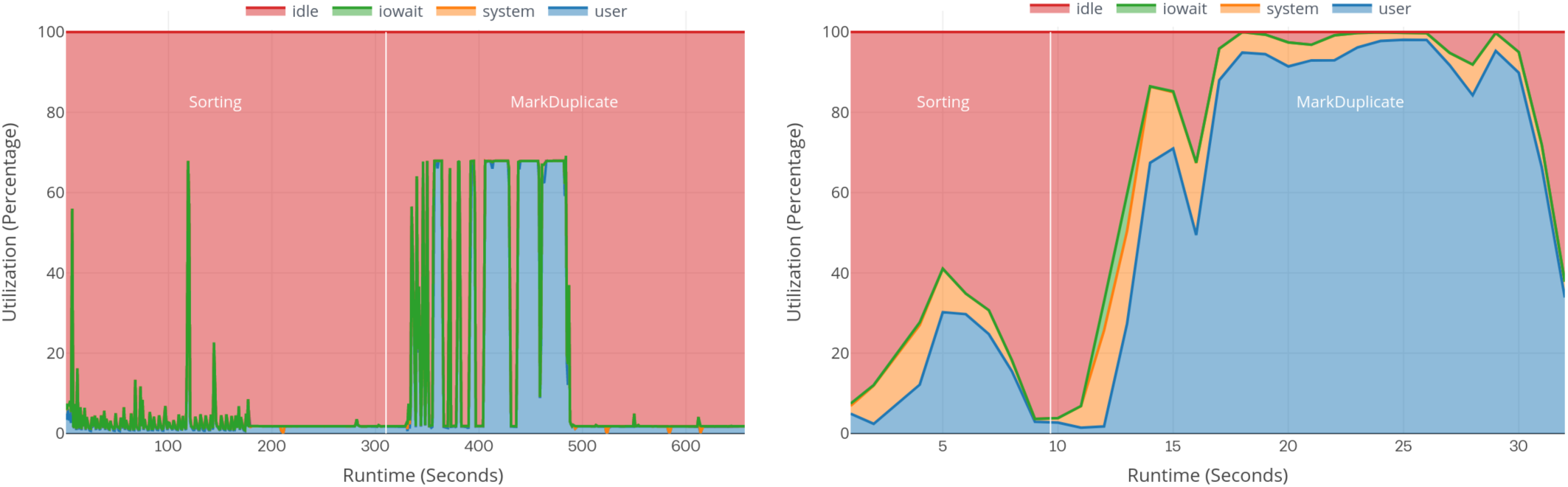
CPU resources utilization for standard Picard (left) as well as ArrowSAM-based (right) sorting and MarkDuplicate for whole exome data.

#### 2) Memory access

`Picard` tools read a whole line from the SAM file and then parse/extract all the fields from it. Using ArrowSAM, we only access those SAM fields which are required by that specific tool to process. In addition, due to the columnar format of ArrowSAM, our implementation is able to better exploit cache locality. Figure 4 shows a comparison of level-1 (L1), level-2 (L2), and last-level cache (LLC) statistics for `Picard` as well as ArrowSAM-based sorting (left) and `MarkDuplicate` (right) applications for whole exome data set. The figure shows that, all levels of cache accesses decrease due to the fewer number of in-memory fields that need to be accessed for the sorting and marking duplicates processes in WES. Cache miss rate also decreases in all cache levels and particularly in L1 cache.

**Fig. 4.**
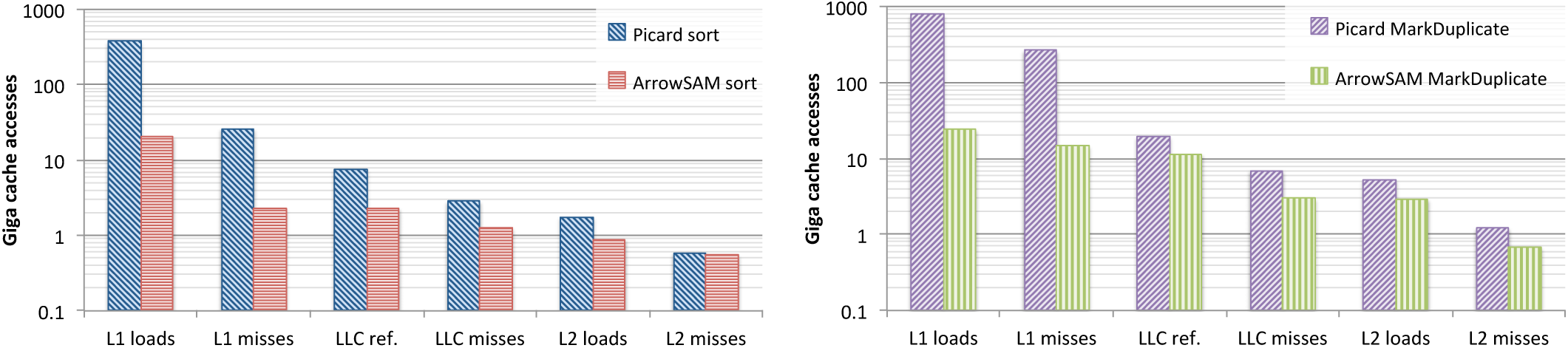
A comparison of level-1 (L1), level-2 (L2), and last-level cache (LLC) statistics for Picard as well as ArrowSAM-based sorting (left) and MarkDuplicate (right) applications for whole exome data set. (LLC ref. stands for LLC references)

#### 3) Memory usage

Unlike other tools, ArrowSAM data resides fully in-memory. Therefore, all the data is placed in a shared memory pool of plasma objects. After sorting, input plasma objects can be removed to free space for the new sorted data which is used in subsequent applications. Other than this, no additional memory is required for intermediate operations. elPrep is an alternative tool that also uses in-memory processing. Memory used by ArrowSAM and elPrep for WES and WGS data sets in pre-processing applications is shown in Table III.

**TABLE III.**
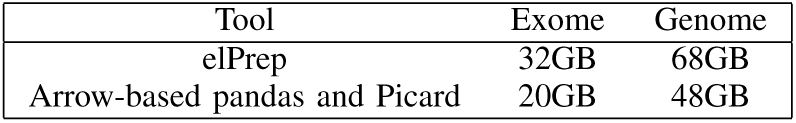
Memory footprint for in-memory processing tools

## VI. Related work

Many in-memory implementations of genomic variant discovery pipelines have been proposed. Almost all these implementations are cluster scaled and do not specifically exploit single node performance. Many use the input data parallelism to distribute the jobs on a cluster [11] and some of them take the benefit of the Apache Spark big data framework for in-memory data management [12], [13]. Our focus is to exploit performance of single node systems.

In addition, some research focuses on creating new genomics tools and algorithms that are more efficient than existing standard genomics pipelines [14]. ADAM [15], a set of formats and APIs uses Apache Avro and Parquet for storage and the Spark programming model for in-memory data caching to reduce the I/O overhead. The results show that ADAM is 3x slower than multi-threaded `Sambamba` in small number of cluster cores up to 64. In `elPrep` [9], the authors report 13x speedup over GATK best practices pipeline for whole-exome and 7.4x faster for whole-genome using maximum memory and storage footprints. The main drawback of these tools is lacking validation in the field which reduces their impact.

Other research focuses on innovative hardware platforms to execute genomics algorithms more efficiently [16]. In [17] a large pool of different types of memories are created and connected to processing resources through the Gen-Z communication protocol to investigate the concept of memory-driven computing. The memory is shared across running processes to avoid intermediate I/O operations. This systems also allows byte-addressability and load/store instructions to access memory. They reported 5.9x speedup on baseline implementation for some assembly algorithms, the source code is not available. Some researchers use high-performance hardware accelerators such as GPUs [18] and FPGAs [19] to accelerate computationally intensive parts of genomics pipelines, but availability of such accelerators in the field remains limited.

## VII. Conclusion

This paper proposed a new in-memory SAM data representation called ArrowSAM that makes use of the columnar in-memory capabilities of Apache Arrow. The paper showed the benefit of using ArrowSAM for genomic data storage and processing for genomics data pre-processing: mapping, sorting and mark duplicates. This allows us to process genomics data in-memory through shared memory plasma objects in parallel without the need for storing intermediate results through I/O into disk. Results show speedup of 28x for sorting and 15x for mark duplicates with respect to I/O based processing, more than 4x and 30% memory access reduction for sorting and mark duplicates, respectively, high CPU resources utilization, as well as better cache locality. These results indicate the potential of adopting a standard in-memory data format and shared memory objects for genomic pipeline processing. Future research will focus on extending our work for the complete genomics variant calling pipeline. In addition, we plan to integrate ArrowSAM into big data frameworks like Apache Spark to enable cluster scale scalability of genomics applications. The code and scripts for running all workflows are freely available at https://github.com/abs-tudelft/ArrowSAM.

## References

[1] D. Gurdasani, M. S. Sandhu, T. Porter, M. O. Pollard, and A. J. Mentzer, “Long reads: their purpose and place,” Human Molecular Genetics, vol. 27, no. R2, pp. R234–R241, 05 2018. [Online]. Available: https://doi.org/10.1093/hmg/ddy177

[2] Y. Diao, A. Roy, and T. Bloom, “Building highly-optimized, low-latency pipelines for genomic data analysis.”

[3] H. Li, “Aligning sequence reads, clone sequences and assembly contigs with bwa-mem,” 2013.

[4] “Picard toolkit,” http://broadinstitute.github.io/picard/, 2019.

[5] A. Tarasov, A. J. Vilella, E. Cuppen, I. J. Nijman, and P. Prins, “Sambamba: fast processing of ngs alignment formats,” Bioinformatics, vol. 31, no. 12, pp. 2032–2034, Jun 2015, 25697820[pmid]. [Online]. Available: https://www.ncbi.nlm.nih.gov/pubmed/25697820

[6] Apache. (2019) Apache arrow: A cross-language development platform for in-memory data. [Online]. Available: https://arrow.apache.org/

[7] ApacheFoundation. (2019) Plasma in-memory object store. [Online]. Available: https://arrow.apache.org/blog/2017/08/08/plasma-in-memory-object-store/

[8] U. L. Technology. (2019) Apache arrow platform. [Online]. Available: https://ursalabs.org/tech/

[9] C. Herzeel, P. Costanza, D. Decap, J. Fostier, and W. Verachtert, “elPrep 4: A multithreaded framework for sequence analysis,” PLOS ONE, vol. 14, no. 2, p. e0209523, Feb. 2019. [Online]. Available: https://doi.org/10.1371/journal.pone.0209523

[10] ApacheFoundation. (2019) Python library for apache arrow. [Online]. Available: https://pypi.org/project/pyarrow/

[11] B. Institute. (2019) Introduction to the gatk best practices. [Online]. Available: https://software.broadinstitute.org/gatk/best-practices/

[12] H. Mushtaq and Z. Al-Ars, “Cluster-based apache spark implementation of the gatk dna analysis pipeline,” in Proceedings of the IEEE International Conference on Bioinformatics and Biomedicine, 2015, pp. 1471–1477. [Online]. Available: https://ieeexplore.ieee.org/abstract/document/7359893

[13] H. Mushtaq, F. Liu, C. Costa, G. Liu, P. Hofstee, and Z. Al-Ars, “Sparkga: A spark framework for cost effective, fast and accurate dna analysis at scale,” in Proceedings of the 8th ACM International Conference on Bioinformatics, Computational Biology,and Health Informatics, ser. ACM-BCB ‘17. New York, NY, USA: ACM, 2017, pp. 148–157. [Online]. Available: http://doi.acm.org/10.1145/3107411.3107438

[14] L. Hasan and Z. Al-Ars, “An efficient and high performance linear recursive variable expansion implementation of the smith-waterman algorithm,” in Proceedings of the IEEE Engineering in Medicine and Biology Conference, 2009, pp. 3845–3848. [Online]. Available: https://ieeexplore.ieee.org/abstract/document/5332567

[15] M. Massie, F. Nothaft, C. Hartl, C. Kozanitis, A. Schumacher, A. D. Joseph, and D. A. Patterson, “ADAM: Genomics formats and processing patterns for cloud scale computing,” UCB/EECS-2013-207, EECS Department, University of California, Berkeley, Tech. Rep., 2013.

[16] L. Hasan and Z. Al-Ars, “An overview of hardware-based acceleration of biological sequence alignment,” in Computational Biology and Applied Bioinformatics. InTech, 2011, pp. 187–202.

[17] M. Becker, M. Chabbi, S. Warnat-Herresthal, K. Klee, J. Schulte-Schrepping, P. Biernat, P. Guenther, K. Bassler, R. Craig, H. Schultze, S. Singhal, T. Ulas, and J. L. Schultze, “Memory-driven computing accelerates genomic data processing,” Jan. 2019. [Online]. Available: https://doi.org/10.1101/519579

[18] E. Houtgast, V. Sima, K. Bertels, and Z. Al-Ars, “Gpu-accelerated bwamem genomic mapping algorithm using adaptive load balancing,” in Architecture of Computing Systems (ARCS). Springer, 2016, pp. 130–142. [Online]. Available: https://link.springer.com/chapter/10.1007/978-3-319-30695-7_10

[19] J. Peltenburg, S. Ren, K. Bertels, and Z. Al-Ars, “Maximizing systolic array efficiency to accelerate the pairhmm forward algorithm,” in IEEE International Conference on Bioinformatics and Biomedicine, 2016, pp. 758–762. [Online]. Available: https://ieeexplore.ieee.org/abstract/document/7822616

